# Cis-inhibition of Notch by Delta controls follicle formation in *Drosophila melanogaster*

**DOI:** 10.1101/2025.03.19.644164

**Authors:** Caroline Vachias, Muriel Grammont

**Affiliations:** CNRS 6293, Clermont University, Inserm U1103, UMR GReD, UFR Médecine, Clermont-Ferrand F-63001, France; Univ Lyon, ENS de Lyon, CNRS UMR 5239, INSERM U1210, Laboratory of Biology and Modeling of the Cell, 15, Parvis Rene Descartes, F-69007, Lyon, France

**Author notes:** Corresponding author: Muriel Grammont.

**Keywords:** cis-inhibition, *Delta*, *Notch*, *Drosophila*, polar cells, follicle formation

## Abstract

Daughters of stem cells often differentiate sequentially in response to inputs from various signalling molecules. We focus on the regulation of Notch signalling in the *Drosophila* germarium. This structure contains several somatic stem cells, which give rise to polar cell or follicular cells precursors. Notch, which is required to form follicles, has been shown to be activated in some stem cells and in the polar cell precursors by a Delta signal produced by the germline cells. However, it is known that removing *Delta* from the germline does not phenocopy Notch phenotype, indicating that the mechanism of N activation and its role remain unclear. Here, we demonstrate that Delta in the soma buffers Notch activity via a cis-inhibition mechanism, resulting in the maintenance of an undifferentiated state. In the somatic stem cells and in the polar cell precursors, somatic Delta prevents Notch from being activated by Delta from the germline or from the soma, respectively. Our data also show that these two steps of Notch activities play a redundant role in follicle formation. Thus, Notch activity depends on both germline and somatic Delta, explaining the phenotype discrepancies between *Notch* and *Delta* ovaries. In addition, we show that somatic Delta is more efficient in initiating pc differentiation than germline Delta. Finally, our work provides a novel example of the importance of the regulation of Notch activity through a cis-inhibitory mechanism.

## Introduction

Stem cells stand out from other cells by their ability to both self-renew and produce differentiated daughter cells. Differentiation of daughter cells is controlled by intrinsic or extrinsic cues. Intrinsic cues often lead to immediate differentiation while differentiation may be delayed when it depends on the spatial influence of the extrinsic cues on the cells. Ovarian follicle formation in *Drosophila* is an ideal system to address this process, as it has been shown that differentiation of the daughter cells of the somatic stem cells does not depend of any lineage, but rather is established progressively through the activities of various signalling pathways (Dai et al., 2017; Melamed et al., 2023; Melamed and Kalderon, 2020a; Nystul and Spradling, 2010a; Reilein et al., 2018, 2017a; Rust and Nystul, 2020; Wang et al., 2021).

A *Drosophila* ovarian follicle is composed of a monolayer of epithelial cells surrounding a cyst of 16 germline cells (fifteen nurse cells and one oocyte). Follicles are continuously produced within the germarium, a structure where both the germline and the somatic stem cells reside (Figure 1A). The germarium is functionally divided into four regions. First, in the anterior part of the germarium (region I), divisions of germline stem cells give rise to cystoblasts that undergo four rounds of incomplete divisions to form 16-cell cysts (King, 1970; Spradling, 1993). Second, in region IIa, these 16-cell cysts interact with a population of about 15 somatic stem cells (ssc). Ssc and their siblings start migrating around germline cysts, that progressively stretch out in between the walls of the germarium (region IIb). The ssc may migrate anteriorly, posteriorly or radially. When moving toward the region IIb, they differentiate as follicular cell precursors (pfc) or polar cell - stalk cell precursors (ppc-sc) (Fadiga and Nystul, 2019; Melamed and Kalderon, 2020a; Nystul and Spradling, 2010b, 2007; Reilein et al., 2017b). Through the region III, the follicle rounds up (at which point it is called a stage 1 follicle) and the ppc-sc become either polar cells (pc), localised at the extremity of each follicle, or stalk cells (sc), which intercalate between adjacent follicles. In parallel, the pfc become the main body follicular cell (mbfc).

**Figure 1.**
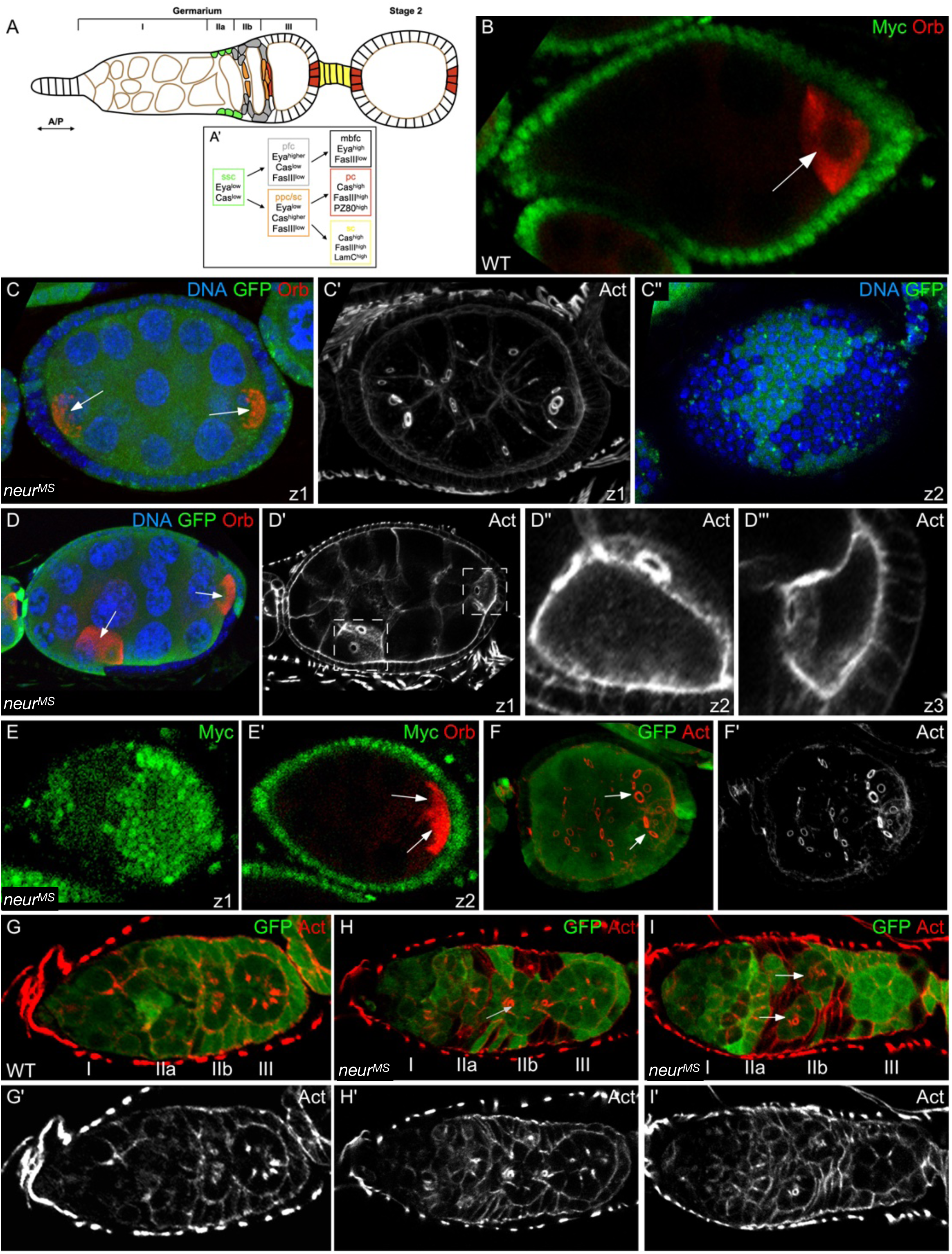
*neur* is required for follicle formation. In all figures, anterior is to the left, mutant clones are marked by the absence of Myc or GFP and roman numerals indicate the different regions of the germarium. (A) Schematic representation of a germarium and a stage 2 follicle and of the different steps of cell differentiation (A’). The arrangement and identities of the different cell types are shown: somatic stem cells (ssc - green), follicular cell precursors (pfc - grey), main body follicular cell (mbfc - black outline), polar cell - stalk cell precursors (ppc-sc - orange), polar cells (pc - red), stalk cells (sc - yellow) and germline cells (brown outline). (B) WT stage 6 follicle. The oocyte, marked by Orb, is localized at the posterior (arrow). (C-F) Stage 6 *neur^MS^* follicles. Each oocyte (Orb, arrows) bear 4 ring canals (marked by Actin). Oocytes are localized either at each extremity (C), or one at the posterior end and one laterally (D) or both at the posterior end (E, F). D’’ and D’’’ are magnified views of the boxed areas in D’. (C, E) Follicles through 2 z-confocal sections (z1 and z2). (G) WT germarium. (H-I) *neur^MS^* germaria. Arrows point to cysts in region IIb that do not display a lens shape and that are not separated from the older (H) or younger (I) cyst.

The somatic stem cell identity is controlled by the opposite activities of the Wnt and the Jak/Stat pathways (Melamed et al., 2023; Melamed and Kalderon, 2020b; Reilein et al., 2017b; Rust and Nystul, 2020). Some ssc migrate radially around the cysts in a Notch dependant manner (Nystul and Spradling, 2010a). When ssc leave the region IIa towards the posterior, their differentiation in pfc or ppc-sc depend on the intrinsic balance between the Eyes Absent (Eya) and Castor (Cas) proteins, themselves being under the control of the Wnt and the Hedgehog signalling pathways (Dai et al., 2017; Reilein et al., 2017a). Eya and Cas are present first at low levels in ssc prior exhibiting stronger levels in the pfc in the case of Eya or to the ppc-sc (in the case of Castor) (Bai and Montell, 2002; Chang et al., 2013; Dai et al., 2017). From the regions IIb and III, the fate of the polar cells and of the stalk cells is progressively established, through the activities of the Notch (N) and Jak/Stat pathways (Assa-Kunik et al., 2007; Grammont and Irvine, 2001; Lopez-Schier and St Johnston, 2001; McGregor et al., 2002). In stage 2 follicles, in addition to *Cas* expression, polar cells start expressing at high levels other specific markers, such as *Fasciclin III* (Fas III), or the PZ80 (in *Fas III*) and A101 (in *neuralized*) reporters, reflecting the last step of differentiation.

During follicle formation, the Notch pathway is required to ssc radial migration, cyst partition and pc differentiation. For all these steps, it has been shown that N is activated by the Delta ligand produced by the germline cells. As the ligand Serrate is not required in the germline or in the soma for any of these processes (Rusconi and Corbin, 1998), removing N in all the somatic cells from region IIa onwards should produce the same phenotypes than removing Dl from the germline. This is in fact largely incorrect. Lack of N leads to an absence of radial migration of the ssc, to the formation of swollen germaria with improperly encapsulated cysts and to a lack of polar cells (Grammont and Irvine, 2001; Lopez-Schier and St Johnston, 2001; Nystul and Spradling, 2010b; Xu et al., 1992). Surprisingly, Dl-germline clones do not lead to swollen germaria, do not affect follicle formation nor the first steps of pc differentiation (Lopez-Schier and St Johnston, 2001; Nystul and Spradling, 2010b; I L Torres et al., 2003). Thus, the mechanism of N activation for cyst enclosure and for pc differentiation remains to be identified. We re-analysed the role of the Dl ligand and of one of its modulators, *neuralized* (*neur*) in these processes. Neur encodes an E3-ubiquitine ligase required for efficient Dl signalling activity through endocytosis (Lai et al., 2001; Pavlopoulos et al., 2001; Yeh et al., 2001).

Our results indicate that *Dl* is required in the soma throughout the germarium to avoid aberrant N activity through a cis-inhibition mechanism and this mechanism is required to control proper timing of pc differentiation. We show that *Dl* somatic expression prevents N from responding to Dl from the germline in region IIa, whereas it prevents N from being activated by the Dl signal from the neighbouring follicular cells in regions IIb/III, restricting the pc fate to the ppc. Our data also show that some ssc and the ppc-sc escape from the cis-inhibition and that these two types of cells play a redundant role in follicle formation.

## Results

### N activation by Dl within the somatic cells are required for follicle enclosure

N activation occurs when ligands and receptor interact in opposite cells (trans-activation), but can be blocked when ligands and receptors are present in the same cell (cis-inhibition). To determine whether Dl activates N between the somatic cells during follicle formation, we analysed the requirement for Neur, which is essential in Dl-expressing cells to activate N in trans. When *neur* clones encompass 25% - 75% of somatic epithelial cells (henceforth referred to *neur^MS^* follicles or *neur^MS^* germaria, Figure S1) follicles containing more than 16 germline cells are frequently observed (78%, n=62). Such compound follicles could result either from abnormal division of germline cells, from abnormal enclosure by the follicular cells in the germarium, or from a collapsing of the stalk after follicle formation, which occurs when Dl is absent from the germline (Isabel L Torres et al., 2003). To distinguish between these possibilities, we first analysed the expression of the Orb oocyte marker and counted the number of ring canals connecting the oocytes. All compound follicles (n=16) contained two Orb-positive oocytes, connected to 4 nurse cells each (Figure 1B-F). Because these oocytes are not always adjacent to each other (they are either one at each end of the follicle (82.7%, Figure 1C); one posterior and one lateral (11.5%, Figure 1D); or both at the posterior end (5.8%, Figure 1E-F), we can rule out that compound follicles are caused by abnormal germline divisions. Compound follicles are thus not due to abnormal division of the germline cells as they always enclose two 16-cell cysts. Next, we investigated whether compound follicles derive from abnormal cyst enclosure in the germarium or from follicle fusion in the vitellarium. Analyses of *neur^MS^* germaria show compound follicles in region III and the presence of round-shaped cysts in region IIb, instead of a single, lens-shaped cyst spreading throughout the depth of the germarium (Compare Figure 1G with 1H and 1I). The separation of the germline cysts by the somatic cells is thus impaired when *neur* somatic cells are present in the germarium. Together, these observations show that *neur* is required in the soma for cyst enclosure and therefore suggest that *Dl* is also required in the soma for this process.

### The germline and soma are redundant sources of Dl for follicle formation

As *neur* is required for the production of active Dl ligands, its requirement for follicle formation suggests that Dl produced by the soma activates N, in parallel to its activation by Dl produced by the germline. We re-examined the tissue-specific requirement of *Dl* during follicle formation by counting the number of compound follicles when either all the germline cells or all the somatic cells (henceforth referred to as *Dl*^G^ or *Dl*^S^ follicles, respectively) were mutant (Figure S1). We found that *Dl*^G^ follicles were always WT (n=12) (Figures 2A, S2A) whereas *Dl*^S^ follicles were occasionally compound (13%, n=23) (Figures 2B, S2B-C). We then analysed ovarioles where both the germline and the soma were mutant for *Dl*. This consistently led to the formation of swollen germaria and huge compound follicles (100%, n=14), a phenotype similar to N mutants (Figure 2C). These data prove that *Dl* is required for follicle formation, and that this role can be fulfilled by either a somatic or a germline source (that we refer to as DlS and DlG, respectively).

**Figure 2.**
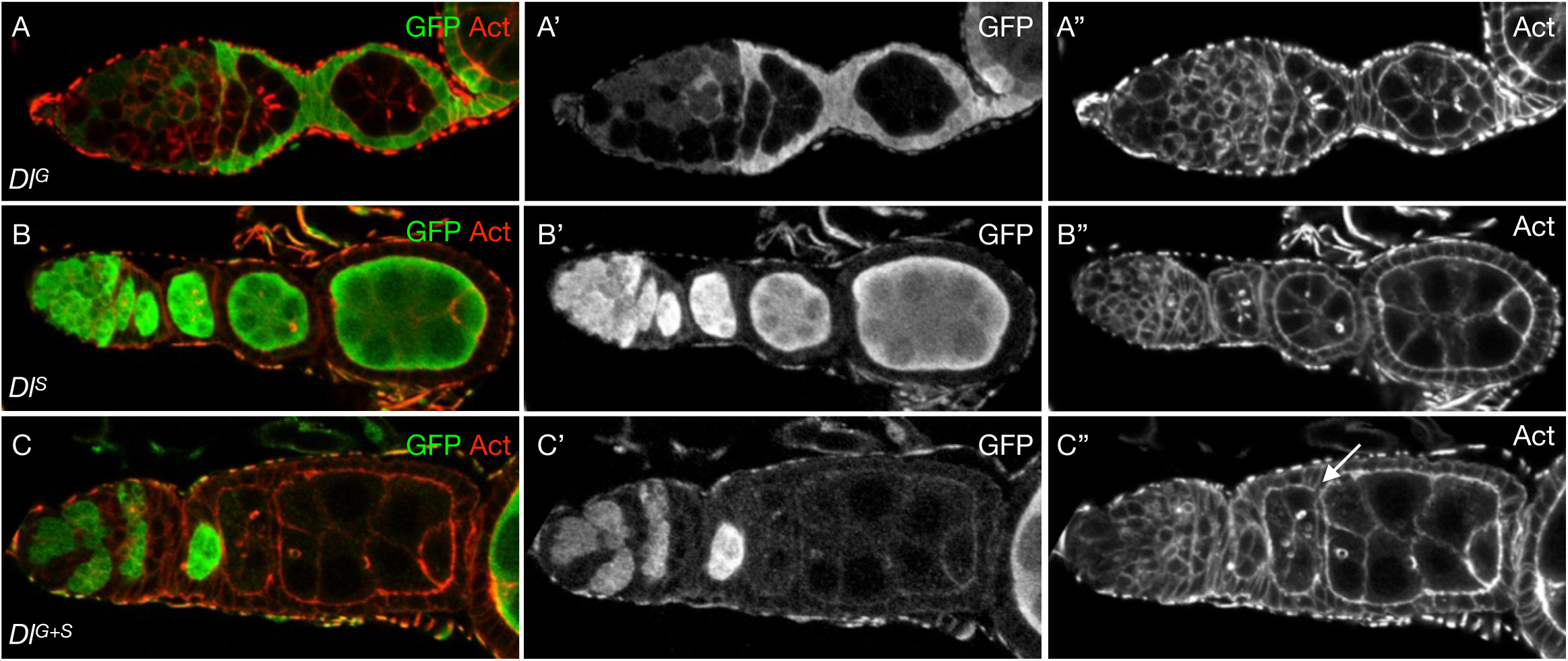
Dl_G_ and Dl_S_ are both required to activate N for follicle formation. Ovarioles with *Dl*-germline (*Dl^G^*) clones (A) or *Dl*-soma (*Dl^S^*) clones (B) or with both germline and soma (*Dl^G+S^*) clones (C). The arrow in C points to adjacent cysts in a compound follicle.

### Cis-interactions between N and DlS downregulate N activity during follicle formation

To further analyse the redundancy between DlS and DlG, we analysed germaria comprising both WT and *Dl* somatic cells (subsequently referred to as *Dl*^MS^; Figure S1). These were frequently compound or fused (74%, n=35) (Figure 3A), revealing that the juxtaposition of somatic mutant cells with WT germline or WT somatic cells is more deleterious for follicle formation than the complete removal of Dl from the germline or from the soma. We hypothesized that this phenotype could come from inappropriate N activation due to the perturbation of a cis-inhibition regulatory mechanism, which has been observed in cells that express both *N* and *Dl* (Del Álamo et al., 2011; Miller et al., 2009). When this mechanism is present, Dl endogenously blocks N and prevents it from being activated by exogenous source of Dl, while in *Dl* cells, N is free to be trans-activated by this source. Because both *Dl* and *N* are expressed in the soma of the ovary, a cis-inhibition mechanism may exist in any somatic cell of the germarium. To test this possibility, we analysed the expression of several reporters for N activity in *Dl*^MS^ germaria (see Materials and Methods). Quantitative analyses have been done with the (P(*GbeSu(H)m8*)) reporter, as it is considered as the most faithful and is therefore the most used since few years (Nystul and Spradling, 2010a; Shi et al., 2026; Troost et al., 2023a, 2023b; Zacharioudaki and Bray, 2014). In WT, we observed that 10 % to 15% of WT germaria expressed the (P(*GbeSu(H)m8*)) reporter in the ssc (region IIa) and in the ppc-sc precursors (region IIb/III) (n=25) (Figure S3). This percentage is similar to what has been observed by Nystul and Spradling (Nystul and Spradling, 2010a), confirming the on-off activity of the N pathway during follicle formation. In contrast, we found that 100% of *Dl*^MS^ germaria displayed N reporter expression in region IIa and 65% in region IIb/III, respectively (Figure 3B, C, n=42). This confirms that inappropriate N signalling occurs in *Dl*^MS^ germaria. In region IIa, N activity was detected in all *Dl* ssc independently of the presence of neighbouring WT somatic cells, whereas N activity in regions IIb/III only occurred in *Dl* somatic cells that are in contact with WT somatic cells (Figure 3B, C). This suggests that N activation depends on DlG in region IIa and on DlS in regions IIb/III. This was confirmed first by examining *Dl*^S^ germaria, which still display multiple cells expressing *GbeSu(H)m8* in region IIa but none in region IIb/III (Figure D, n=12). Second, we analysed N reporter expression in germaria with somatic *Dl* clones and *Dl* germline cysts (that we refer to as *Dl*^G+MS^; Figure S1). No expression of the reporter was observed in region IIa while expression is still detected when *Dl* somatic cells were in contact with WT somatic cells in region IIb/III (Figure 3E, n=8). This confirms that in *Dl* somatic cells, N is activated by DlG in region IIa, but by DlS in region IIb/III. Altogether, these data show that all the somatic cells in the germarium express *Dl* and that this expression is crucial to prevent N from responding to DlG in region IIa, and to DlS in region IIb/III. These data also demonstrate that the DlG is unable to activate N reporter in any somatic cells in regions IIb and III of the germarium.

**Figure 3.**
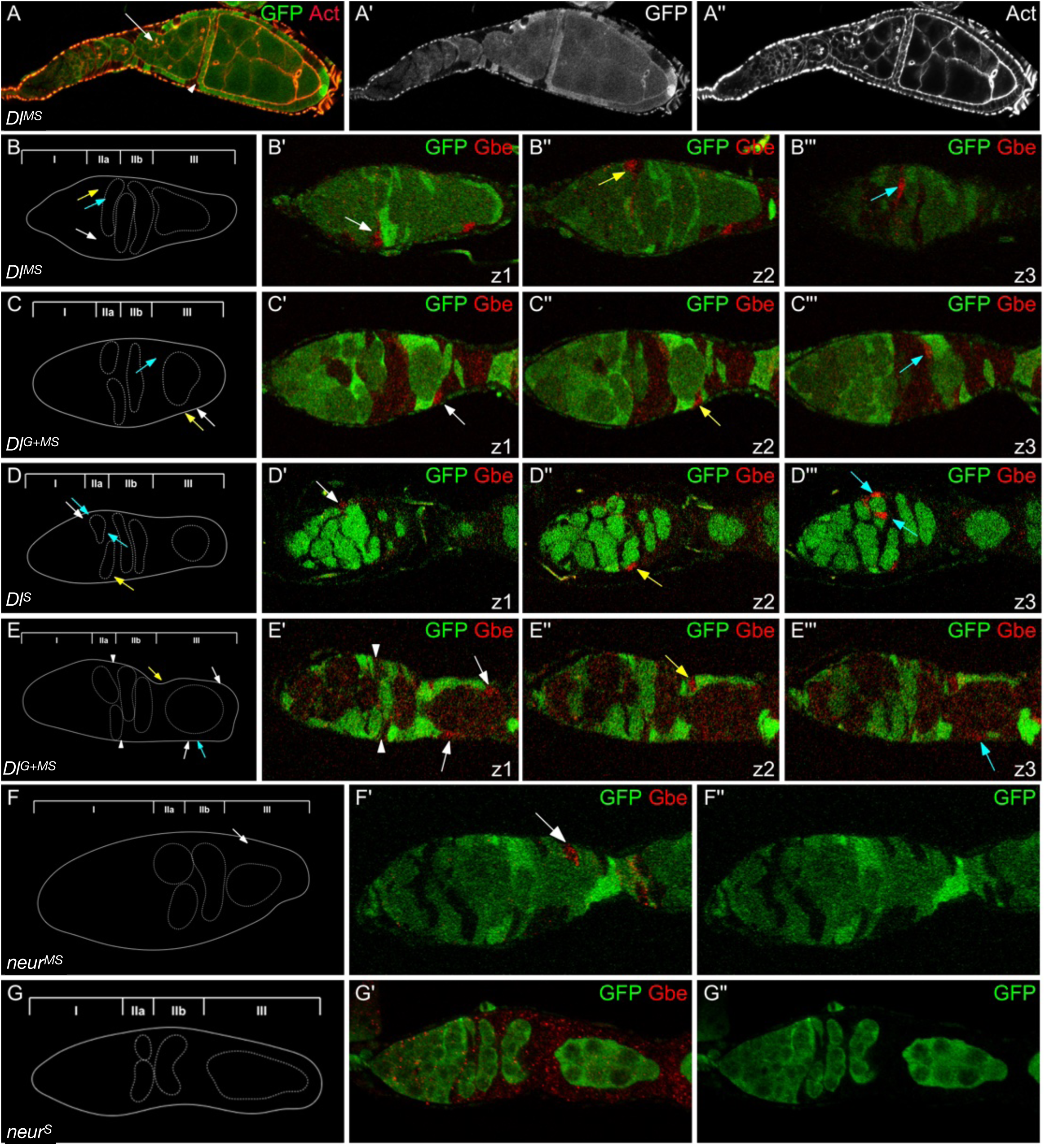
Dl_S_ cis-inhibits N throughout the germarium. (A) *Dl^MS^* ovariole with compound (arrow) and fused follicles (arrowhead). (B-G) Germaria from females carrying the *GbeSu(H)m8* reporter, through three representative z-sections (z1 to z3). A schematic of each germaria is presented on the left. The position of each germline cyst (white dotted line) in regions IIa, IIb and III is shown. Arrows indicates the *GbeSu(H)m8-*expressing cells in different z sections (white for z1, yellow for z2 and blue for z3). (B) *Dl^MS^* germarium with three *GbeSu(H)m8* cells (arrows) in region IIa. (C) *Dl^G+MS^* germarium. Three *GbeSu(H)m8* MBFC (arrows) are visible in region III. All cysts are WT except the *Dl* cyst in region IIb. (D) *Dl^S^* germarium. Four *GbeSu(H)m8* cells (arrows) are visible in region IIa. (E) *Dl^G+MS^* germarium. *Dl* cysts are present throughout the germarium. No *Dl* cells (arrowheads) in region IIa express *GbeSu(H)m8* whereas at least four mbfc expressing *GbeSu(H)m8* are present in region III (arrows). (F) *neur^MS^* germarium with mbfc expressing *GbeSu(H)m8* (arrow) in region III. (G) *neur^S^*germarium. No cells expressed *GbeSu(H)m8* throughout the germarium.

We next analysed N reporter expression in *neur*^MS^ germaria. We did not detect any N reporter expression in the ssc in region IIa, in contrast to region IIb/III, where expression is detected in the *neur* somatic cells that are in contact with WT somatic cells (Figure 3F, n=15). Similar to *Dl^S^* germaria, this expression is not detected *neur^S^ germaria* (Figure 3G, n=6). Thus, this activation depends on the signal produced by neighbouring WT cells. From these experiments, we conclude that DlS activates N in region III in a *neur*-dependant manner when the cis-inhibition mechanism is impaired.

### DlS prevents premature and ectopic polar cell differentiation

Since strong N activity correlates with polar cell fate induction (Grammont and Irvine, 2001; Lopez-Schier and St Johnston, 2001; Nystul and Spradling, 2010b), we then asked whether too many cells adopt a polar cell fate in a *Dl*^MS^ germaria by looking at Castor and PZ80 expression. In such germaria, Cas was strongly expressed in *Dl s*sc in region IIa (Figure 4A, B, n=8). In these cells, the fluorescence intensity was about four times more than those measured in WT ssc (Figure S4A, S4B) and correlated with high levels of N reporter expression (Figure 4C, n=5). Our data also demonstrate that this strong Cas level depends on N activity, as it is lost in *Dl*^G^ germarium (Figures 4D, S4D, n=6), demonstrating that the cis-inhibition mechanism prevent ssc to prematurely express a polar cell marker.

**Figure 4.**
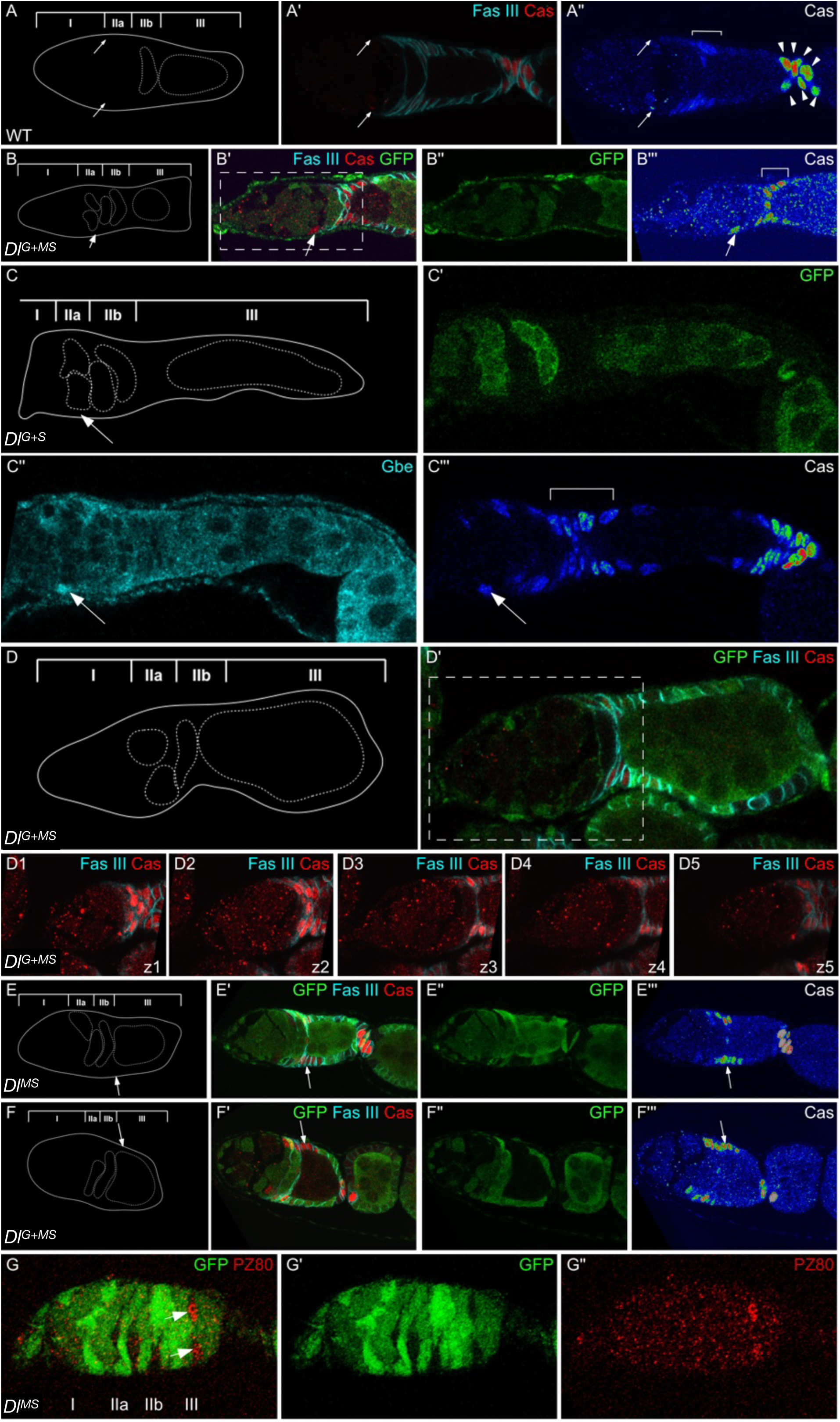
DlS controls polar cell marker expression in the germarium. The use of a color gradient allows the visualization of differences in intensities of accumulation from none (blue) to strongest (red) (A’’, B’’’, C’’’, E’’’, F’’’). (A-F) A schematic of each germaria is presented on the left. The position of each cyst (white dotted line) in regions IIa, IIb and III is shown. (A) WT germarium showing the different levels of Cas expression in the ssc and pfc (weak level, arrows), the ppc-sc (intermediate level, brackets), the pc and the sc (high level, arrowhead). (B) *Dl^G+S^* germarium. A *Dl* cyst is present in region III. All the somatic cells are mutant. The arrow points to a stem cell expressing high levels of Castor. High Cas expression is also seen in the pfc and ppc-sc (brackets) at the border between the regions IIb and III. (C) *Dl^G+MS^* germarium from a female carrying the *GbeSu(H)m8* reporter. The arrow points to cells expressing intermediate levels of Castor and high levels of N activity in region IIa. High Cas expression is also seen in pfc and ppc-sc precursors. A *Dl* cyst is present in region IIb. (D) *Dl^G+MS^* germarium through 5 consecutive z-confocal sections (z1 to z5). No cells are expressing Cas before region IIb. All the cysts present in regions IIa and IIb are mutant for *Dl*. (E) *Dl^MS^* germarium. The arrow points to mbfc expressing high levels of Castor in region III. (F) *Dl^G+MS^* germarium. The arrow points to mbfc expressing high levels of Castor in region III. A *Dl* cyst is present in region III. (G) *Dl^MS^* germarium from a female carrying the PZ80 enhancer-trap. The arrows point to mbfc expressing high levels of β-gal in region III.

In region IIb/III of *Dl*^MS^ germarium, Cas was strongly expressed in *Dl* cells that are in between adjacent cysts, which are considered to be the ppc-sc (Figure 4A-B, E-F, n=24). The fluorescence intensity in *Dl* ppc-sc was about twice that of WT pc (Figure S4A and S4C). These results indicate that DlS is required to prevent high Cas expression in the ppc-sc. This was observed independently of the genotype of the germline (Figure 4B, D, F) and of the somatic cells (Figure 4C). *Dl* mbfc also exhibit a strong Cas level, if they are not too far from the anterior extremity of the follicle (Figure 4B, 4C, 4E, 4F and Figure S4D, S4E). No correlation was observed between high level of Cas expression and of N activity in the mbfc, indicating that Cas up-regulation is not driven by increased N signalling (Figure S4E, n=6). This is consistent with previous results that showed that Cas expression is not modified in *N* cells (Chang et al., 2013). Upregulation of Cas expression was less detectable in *Dl* mbfc close to the end of germarium, and was completely absent from stage 2 onwards (Figure S4E, n=24). These results indicate that DlS helps to keep Castor at low level in the pfc/mbfc that are close to region IIb.

Finally, we analysed PZ80 expression, which is considered to be a late marker of polar cell fate whose expression usually starting only from stage 2 follicles onwards. In *Dl*^MS^ germarium, while no expression was detected in regions IIa and IIb, we did observe premature PZ80 expression in *Dl* mbfc when they were in contact with WT mbfc in region III (Figure 4G, n=5). This indicates that DlS prevents polar cell differentiation in the MBFC. To conclude, DlS is required to maintain an intermediate Cas level in the ppc-sc and prevents the mbfc from expressing both Cas and PZ80. Altogether, we can conclude that Dls prevents most of the somatic cells from acquiring prematurely a polar cell fate during cyst enclosure, supporting the possibility that the cis-interaction between N and Dl is important to maintain an undifferentiated state.

### Robust polar cell differentiation depends sequentially on DlS and DlG production

The literature showed that DlG is required to induce polar cell fate (Lopez-Schier and St Johnston, 2001; I L Torres et al., 2003). This conclusion was reached by examining the effect of *Dl*-germline clones without accounting for the presence of *Dl*^SM^ clones. As our data indicate that polar cell differentiation is impaired in *Dl*^SM^ germaria, we decided to re-examine polar cell differentiation in *Dl*^G^ or in *Dl*^S^ follicles. In WT, the vast majority of stage 2 follicles (90-100 %) display polar cell marker expression at the anterior and posterior ends. We observe that both *Dl*^G^ or *Dl*^S^ follicles display delays in the appearance of polar cell markers in the anterior and posterior (Figure 5A), but that polar cell markers are detected in absence of DlG or of DlS (Figures 5B and 5C, S5A). Delays are more severe when *Dl* is removed from the soma than when *Dl* is removed from the germline. Notably, polar cell differentiation rarely occurs at the posterior of *Dl*^S^ follicles prior to stage 4. However, DlG is required and sufficient to maintain this fate after stage 4. This is in agreement with what has been previously described: *Dl^G^* follicles fuse with anterior ones and the expression of some polar cell-specific markers, such as PZ80, become undetectable (Figure S5B). We also analysed the expression of the N activity reporter in young *Dl*^G^ or *Dl*^S^ follicles. We have previously shown that the polar cell pair formation is a two-step N-dependent mechanism, with first the selection of one pc, which then activates N in one ppc (Vachias et al., 2010). This activation prevents this ppc to undergo apoptosis. As expected, most of the *Dl*^S^ follicles did not show *GbeSu(H)m8* expression before stage 4 (compare Figure 5D and 5E, n=16), confirming that DlS production is required to turn on N reporter expression during polar cell selection. *Dl*^G^ follicles often showed expression at both poles at stages 3 or 4, but the expression was weaker than in WT (Figure 5F, n=16), indicating that DlG contributes to some extend to the activation of the N reporter before stage 4. Altogether, these data show that Dl must be produced in both tissues to ensure robust polar cell formation, with DlS playing a preponderant role from stages 1 to 3 and DlG being essential to maintain this fate from stage 4 onwards.

**Figure 5.**
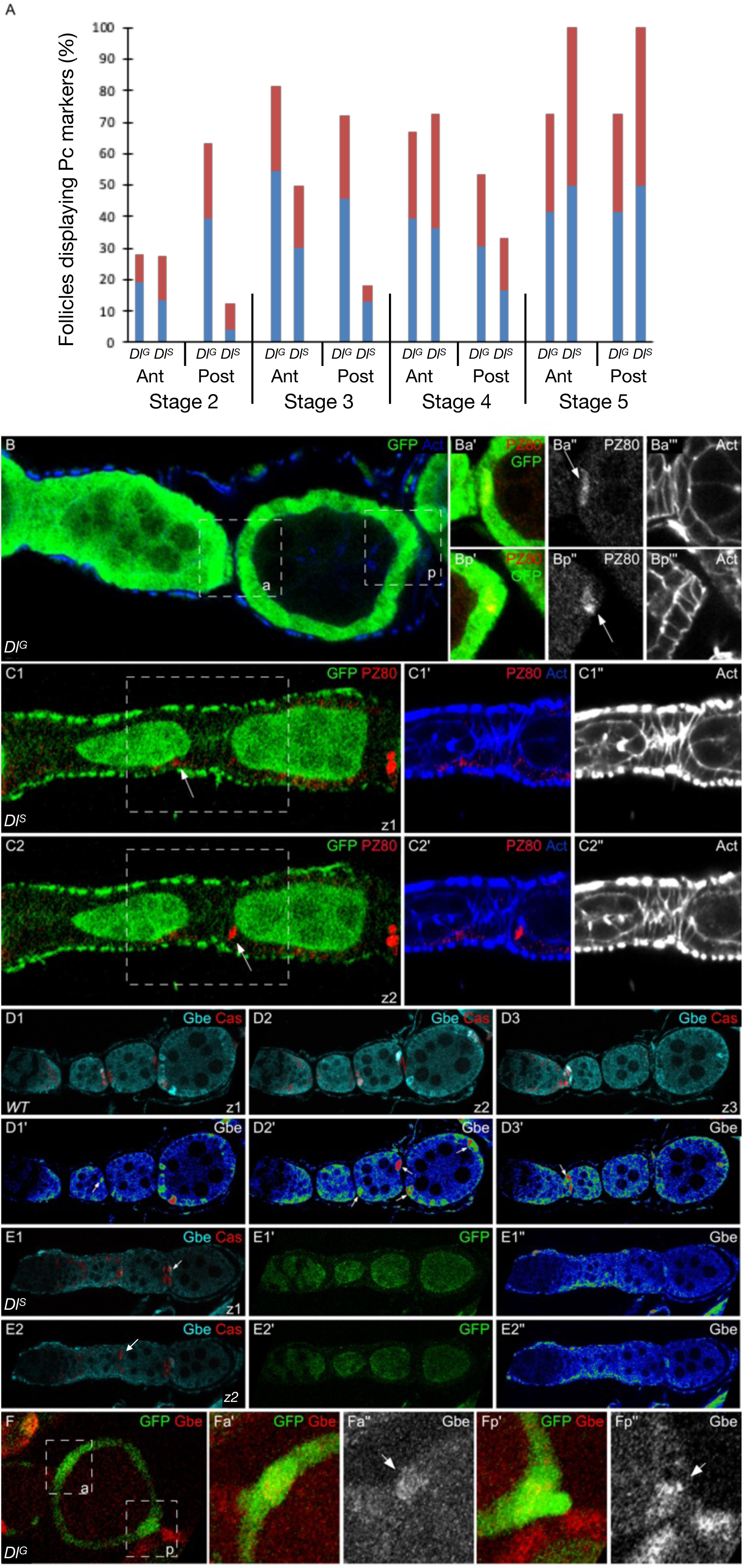
Robust polar cell differentiation requires DlG and DlS. (A) Percentages of follicles containing one or more cells expressing either PZ80 (blue) or Cas (red) polar cell markers in *Dl^G^* follicles or in *Dl*^S^ follicles from stages 2 to 5. Over 14 clusters were counted per stage for each type of *Dl* follicles. (B) Stage 3 *Dl^G^* follicle from a female carrying the PZ80 enhancer-trap. One anterior and two posterior cells express PZ80 (arrow). Ba’ - Ba’’’ and Bp - Bp’’’ are magnified views of the anterior (Ba) and posterior (Bp) boxed areas in B. (C) Stage 2 and 3 *Dl^S^* follicles from a female carrying the PZ80 enhancer-trap, through 2 consecutive z-confocal sections (z1 and z2). One posterior cell of the stage 2 follicle and one anterior cell (arrows) of the stage 3 follicle display PZ80 expression. C1’-C1’’ and C2’-C2’’ are magnified views of the boxed areas in C1 and C2, respectively. (D) WT ovariole from a female carrying the *GbeSu(H)m8* reporter through 3 consecutive z-confocal sections (z1 to z3). The arrows point to polar cells expressing *GbeSu(H)m8*. (E) *Dl^S^* ovariole from a female carrying the *GbeSu(H)m8* reporter through 2 consecutive z-confocal sections (z1 to z2). No expression is detected up to the anterior of stage 4 follicle (arrow). (F) Stage 3 *Dl^G^* follicle from a female carrying the *GbeSu(H)m8* reporter. N activity is detected at the anterior and posterior polar cells (arrows). Fa’, Fa’’, Fp’, Fp’’ are magnified views of the anterior (a) and posterior (p) boxed areas in F.

## Discussion

Previous analyses have shown that N is required for the radial migration of ssc and their siblings in region IIa of the germarium and for the differentiation of the ppc starting in region IIb (Grammont and Irvine, 2001; Lopez-Schier and St Johnston, 2001; Nystul and Spradling, 2010b). Though both processes are known to depend on DlG production, the discrepancies in *N*^S^ and *Dl*^G^ phenotypes implied that the current understanding of the activation of this pathway was incorrect (Lopez-Schier and St Johnston, 2001; Nystul and Spradling, 2010b; I L Torres et al., 2003). By revisiting this statement, our study now demonstrates that both Dl production sites, the germline and the soma, act in a complementary manner to ensure cyst production. Moreover, our data also reveal that DlS is required to buffer N against unintended activation in all the somatic cells from region IIa to region III of the germarium. In region IIa, DlS prevents N from responding to DlG, whereas in region IIb/III, DlS prevents pfc and ppc-sc from responding to DlS produced by the neighbouring mbfc (Figure 6). This regulation prevents premature and ectopic expression of polar cell markers. Finally, we establish that timely pc differentiation depends on N activation by DlS in the germarium and young stages while pc fate maintenance requires DlG from stage 4 onwards.

**Figure 6.**
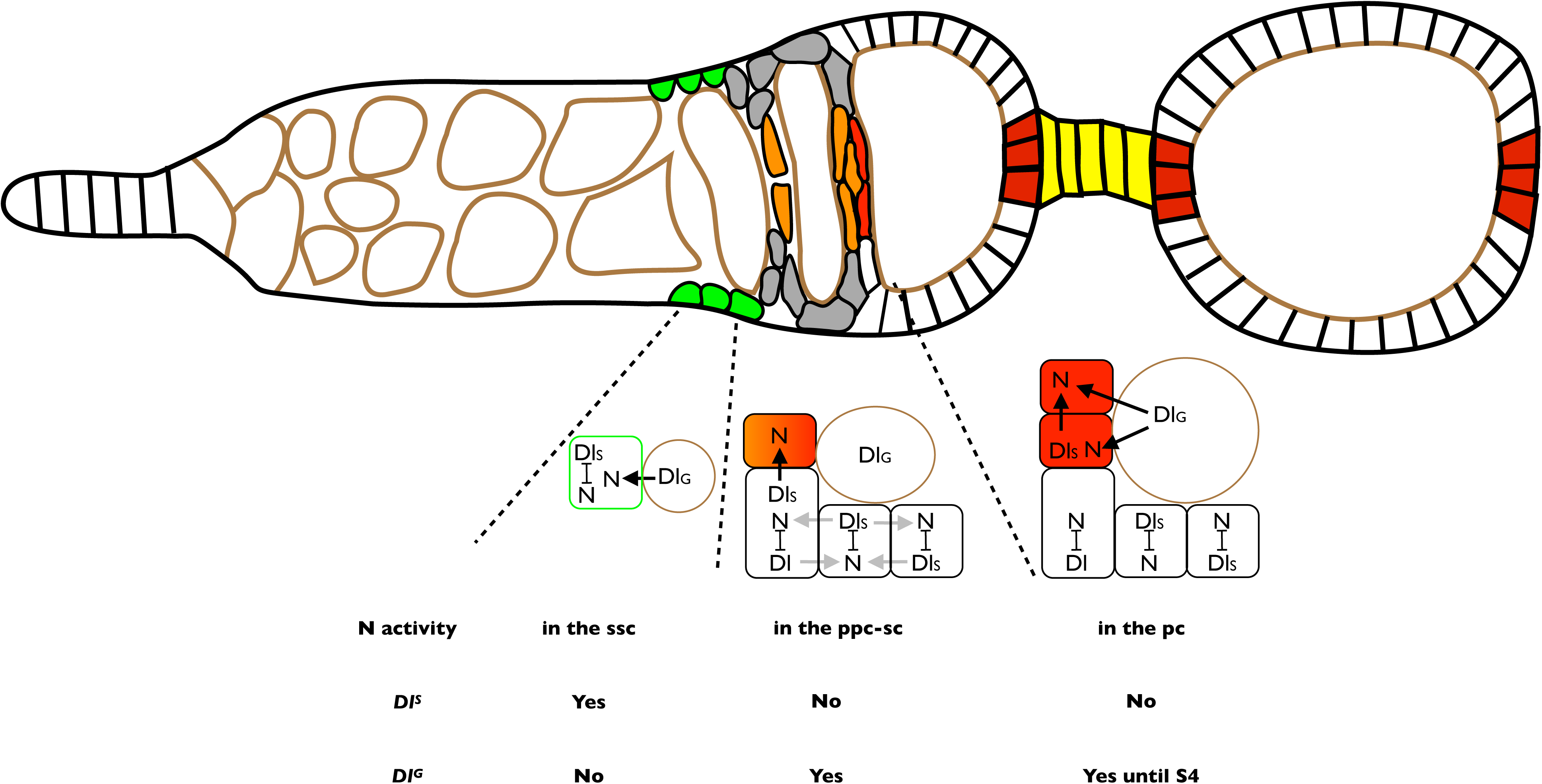
N activity during follicle formation. Schematic representation of a germarium and a stage 2 follicle with the different steps of cell differentiation in WT. The arrangement and identities of the different cell types are as in Figure 1A. In region IIa of WT follicles, DlS cis-inhibits N in all the cells to prevent N to be highly activated by DlG (black arrow). In region IIb/III, DlS cis-inhibits N to prevent it to be activated from the neighboring somatic cells (light grey arrow). DlG is unable to promote activation. Only the pc-sc precursors activate N, possibly due to the activity of Fringe in these cells, which renders the cells more efficient to respond to Dl. In young follicles, DlS and DlG promote N activation on the polar cells (black arrows) DlS still inhibits N in the mbfc. In *Dl*^S^ follicles: N activity is detected in many ssc, but not in the pc-sc precursors. In the polar cells, N activation is not detected. In *Dl*^G^ follicles: N activity is not detected in the ssc, but is activated in the pc-sc precursors. In the polar cells, N activation is detected until stage 4.

Evidence for a cis-inhibitory mechanism during *Drosophila* oogenesis has already been shown in mbfc between stages 2 and 6 (Palmer et al., 2014; Poulton et al., 2011), where DlS blocks N from a ligand-independent activation (Deng et al., 2001). During follicle formation, our data show that the N reporter is not active in the absence of both DlG and DlS, implying that ligand-independent N activation does not occur. Thus, the cis-interactions between Dl^S^ and N permanently present in the germarium must be overcome to allow N activation and suggests that regulators must be present to counteract the cis-inhibition mechanism. One of them is likely to be Fringe, which is known to facilitate N activation when *Dl* is expressed by the neighbouring cells (Grammont and Irvine, 2001; Moloney et al., 2000). *fringe* expression in the ppc likely allows N to be activated in these cells by DlS from the neighbouring somatic cells, despite the cis-inhibition mechanism present in the ppc. However, it is unlikely that *fringe* plays a role to help the ssc to respond to DlG, due to its expression pattern. One possibility could come from the localisation of N receptor in the ssc. As these cells are not epithelial cells yet, transmembrane protein trafficking could allow some N receptors to escape from a cis-interaction with Dl and to be present at the interface with the germline. In parallel, we found that DlG is unable to activate N in any *Dl* somatic cells from region IIb. This could also be due to the transmembrane protein trafficking in epithelial cell, leading to the absence of N at the apical surface.

Several analyses have established the crucial role of Notch for cyst enclosure. This process begins with the radial migration of ssc and their siblings around the germline cysts, followed by the differentiation of some of them in ppc-sc (Fadiga and Nystul, 2019; Melamed and Kalderon, 2020a; Nystul and Spradling, 2010b, 2007; Reilein et al., 2017b). Our results reveal a redundant N function during this process (Figure 6). Indeed, in absence of Dls, follicles are produced and N activity is detected only in the ssc and their siblings of region IIa. This activity is thus sufficient to assure follicle formation. In the absence of DlG, the only cells that display N activity are the ppc-sc and the pc, implying that these cells are also sufficient to promote proper cyst enclosure and follicle individualisation. Altogether, our data support a functional link between the ssc and pc-sc precursors. We propose that the N activation in ssc and ppc-sc assign to these cells the ability to form a follicle. This leads to the intriguing question about whether the N-positive ssc could be the first sign of ppc-sc. This temporary N activity in region IIa may start be the starting point of the polar cell program. Since only 10% to 15% of the ssc display a N activity in fixed tissues, monitoring the dynamics of this activation would be informative to determine whether all ssc get activated at one point or only few of them do. If too many ssc or siblings get activated, it will be unlikely that those represent the pool of cells that will likely give rise to ppc-sc. Recent data also demonstrate that follicle formation is dependent on the germline cell contractility and the somatic cell adhesive properties (Chanet and Huynh, 2020). Analyses of these features in the genetic conditions used in this study would likely be instructive to understand the differences in terms of phenotypes.

The literature suggests that the role of the Hedgehog pathway on Cas and Eya expression is responsible for initiating follicular cell fate specification (Bai and Montell, 2002; Chang et al., 2013). One important aspect of their work was to show the repression exerted by Eya on Cas, which leads to mutually exclusive expression patterns. However, it is unclear why both proteins can be detected at low levels in the cells that reside in region IIa. Our results now show that DlS is required for downregulating Cas in these cells, which implies that both Notch and Hedgehog signalling work together to prevent early follicle cell fate specification, and to allow the maintenance of an undifferentiated state. This is supported by our observation that shows that the juxtaposition of somatic cells with different levels of Dl is more deleterious to follicle formation than the complete absence of Dl would be. This juxtaposition leads to the expression of polar cell markers in the MBFC. Maintenance of an undifferentiated state is likely to be the main role of the cis-inhibition mechanism present in these cells and is crucial for cyst formation.

Our results highlight the importance of protecting cells from inappropriate N activation by signals that may originate in surrounding cells. Although only a few cases of developmental processes using cis-regulation of N by endogenous Dl have been uncovered so far in both vertebrates and invertebrates (Becam et al., 2010; Del Álamo et al., 2011; Miller et al., 2009), this work points to the conclusion that this mechanism is likely to be more common than expected.

## Materials and methods

### Drosophila Stocks and Crosses

CantonS is used as WT. *neur^if65^* and *Dl^rev10^* are null alleles (de Celis et al., 1993; de-la-Concha et al., 1988; Heitzler and Simpson, 1991; Sun and Artavanis-Tsakonas, 1996). The Notch activity reporter line used is GbeSu(H)m8-lacZ (Furriols and Bray, 2001). Fly stocks were cultured at 25°C on standard food. Clones were generated by Flipase-mediated mitotic recombination on FRT82D chromosomes carrying either GFP or Myc as markers (Golic and Lindquist, 1989; Xu and Rubin, 1993). Flipase expression was induced by heat shocking two-day old females at 38°C for 1 hour, then the females were placed at 25°C and were dissected 9 to 12 days after the heat shock. Myc expression was induced by heat shocking females at 37°C for 1 h, four hours before dissection.

### Follicle Staining

Pc identity was assayed by the expression of the FasIII protein and of the PZ80 enhancer trap, inserted in the *fasIII* gene, (Karpen and Spradling, 1992), which are specifically expressed in the pc from stage 1 onwards. The A101 marker (an enhancer-trap within *neur*) was not used in order to prevent feedback from affecting marker expression when the N pathway was manipulated. The following primary antibodies were used: goat anti-ß-galactosidase (1:1000, Biogenesis), rabbit anti-ß-galactosidase (1:2000, Cappel), rabbit anti-Myc (1:100, Santa Cruz), mouse anti-GFP (1:500, Sigma-Aldrich), goat anti-GFP (1:1000, AbCam), mouse anti-FasIII 7G10 (1:200, DSHB), Rabbit anti-Castor (1:5000; a gift from Dr Ward D. Odenwald) and mouse anti-orb 6H4 and 4H8 (1:100, DSHB). Actin staining is realized by using Rhodamin-Phalloidin (Molecular Probes). Secondary antibodies coupled with Cy5, Cy3 (Jackson Immunoresearch) or Alexa Fluor 488 (Molecular Probes) are used at 1/1000.

Ovaries from females were dissected directly into fixative 3 to 4 days after Flipase induction and stained following the protocol described in (Grammont and Irvine, 2001). To avoid fluctuations of the depth of the follicles that are squeezed by the coverslip, each slide contains 15 ovaries, from which S11 to S14 are removed. After dissection of the follicles, most of the PBS is removed and 20µl of the Imaging medium (PBS/Glycerol (25/75) (v/v)) is added before being covered by a 22/32 mm coverslip.

### Confocal Microscopy

Preparations were examined using confocal microscope (LSM 710 and LSM 700; Carl Zeiss MicroImagin, Inc.) with 40x/NA 1.3 plan-Neofluar and 63x/NA 1.4 plan-Apochromat. Imaging was performed at RT. Images were examined using ImageJ (http://imagej.nih.gov/ij/).

### Statistical analysis

All data graphs are reported as mean ± standard deviation. Statistical analysis was performed using R software (RStudio, Inc), with alpha levels for all statistical tests set to 5%.

## Acknowledgments

We thank DSHB, Bloomington Stock Center; Lyon Bio Image and Arthrotools of the SFR Bioscience (UMS3444/US8) for flies and reagents, Dr W. Odenwald for reagents, P. Das for critical comments on the manuscript. This work received financial support by the Agence Nationale de la recherche (ANR-22-CE13-0017-01, ForcesOnCell to MG) (https://anr.fr), the Centre National pour la Recherche Scientifique (https://www.cnrs.fr), and the Ecole Normale Supérieure of Lyon (http://www.ens-lyon.fr). The funders had no role in study design, data collection and analysis, decision to publish, or preparation of the manuscript.

## Author Contributions

Conceptualization: Caroline Vachias, Muriel Grammont.

Data curation: Caroline Vachias, Muriel Grammont.

Formal analysis: Caroline Vachias, Muriel Grammont.

Funding acquisition: Muriel Grammont.

Investigation: Caroline Vachias, Muriel Grammont.

Methodology: Caroline Vachias, Muriel Grammont.

Project administration: Caroline Vachias, Muriel Grammont.

Supervision: Muriel Grammont.

Validation: Caroline Vachias, Muriel Grammont.

Visualization: Caroline Vachias, Muriel Grammont.

Writing – original draft: Caroline Vachias, Muriel Grammont.

Writing – review & editing: Muriel Grammont.

---

**Figure S1.**
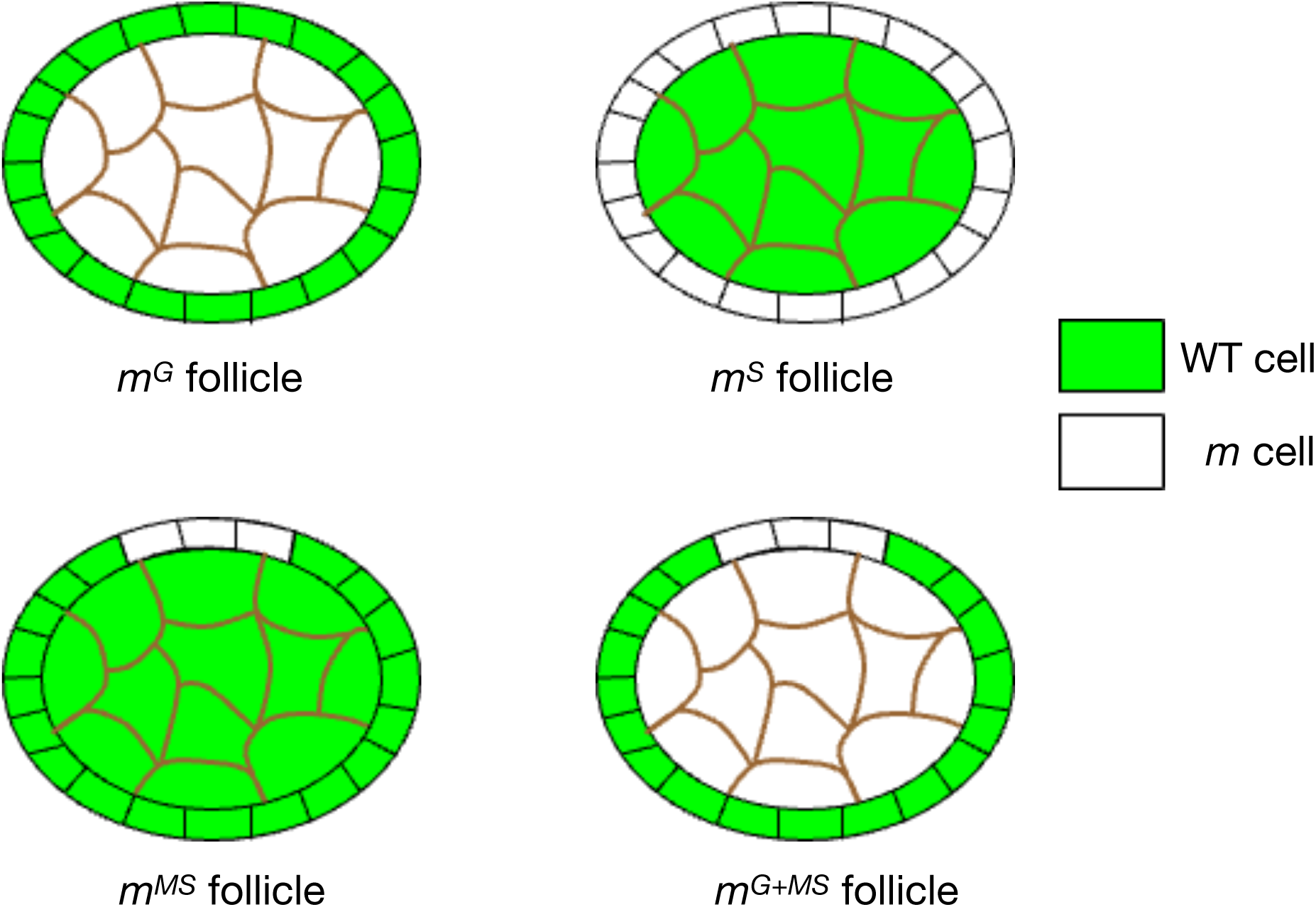
The different types of mutant follicles or germaria. Brown and black outlines correspond to germline and somatic cells, respectively. (A) A *m^G^* follicle is composed of a cyst mutant for either *Dl* or *neur* surrounded by WT somatic cells. A *m*^G^ germarium contains at least one mutant cyst. (B) A *m^S^* follicle is composed of a WT cyst surrounded by somatic cells mutant for *Dl* or *neur*. A *m^S^* germarium contains WT cysts and *Dl* or *neur* somatic cells. (C) A *m^MS^* follicle is composed of a WT cyst surrounded by mosaic somatic cells for *Dl* or *neur*. A *m^MS^* germarium contains WT cysts and mosaic somatic cells. (D) A *m^G+MS^* follicle is composed of a mutant cyst surrounded by mosaic somatic cells. A *m^G+MS^*germarium contains at least one mutant cyst and mosaic somatic cells.

**Figure S2.**
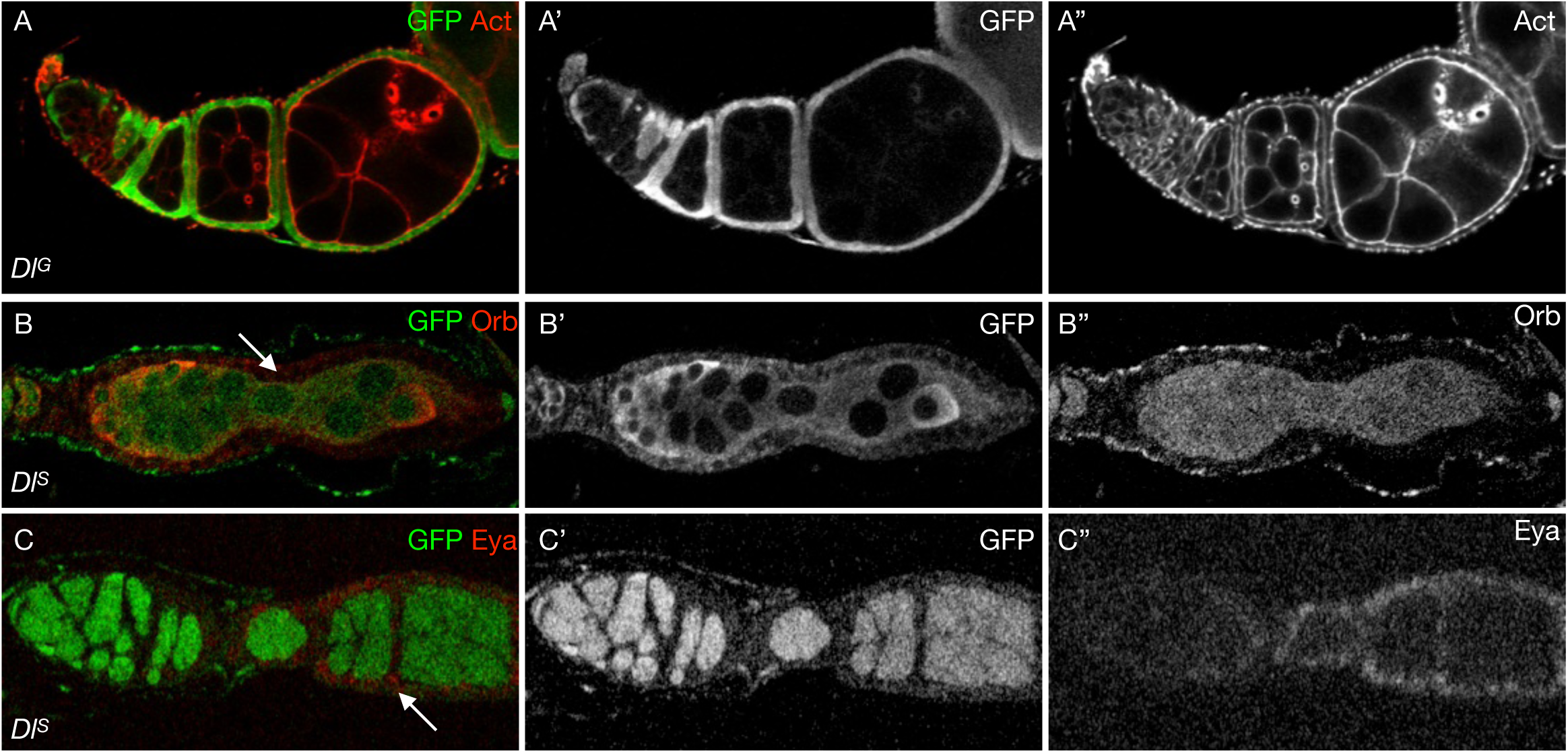
DlG and DlS are both required to activate N for follicle formation. Ovarioles with *Dl*-germline (*Dl^G^*) clones (A) or *Dl*-soma (*Dl^S^*) follicles (B, C). The arrows point to compound (B) or fused follicles (C).

**Figure S3.**
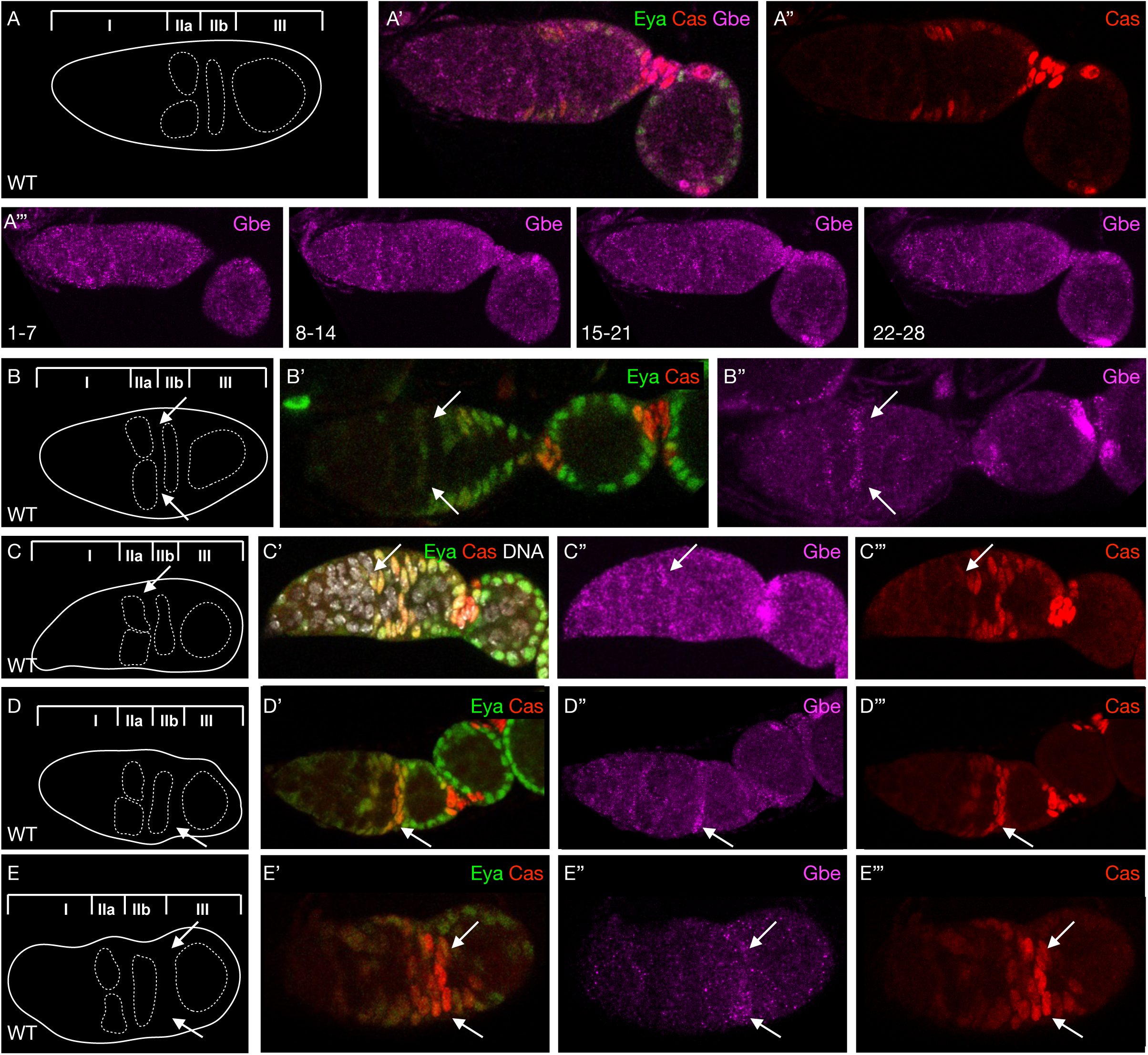
*GbeSu(H)m8* positive cells localization in WT germaria. (A-E) Germaria from females carrying the *GbeSu(H)m8* reporter. A schematic of each germaria is presented on the left. The position of each germline cyst (white dotted line) in regions IIa, IIb and III is shown. Arrows indicates the *GbeSu(H)m8-*expressing cells. (A) Germarium with no *GbeSu(H)m8* positive cells. To appreciate the absence of *GbeSu(H)m8* positive cells, about 6 consecutive z-sections were merged, throughout the depth of the germarium. (B) Germarium with two *GbeSu(H)m8* positive cell in region IIa (arrows). (C) Germarium with one *GbeSu(H)m8* positive cell in region IIa (arrow). (D) Germarium with one *GbeSu(H)m8* positive cell at the border between region IIb and III (arrow). (E) Germarium with three *GbeSu(H)m8* positive cells at the border between region IIb and III (arrow).

**Figure S4.**
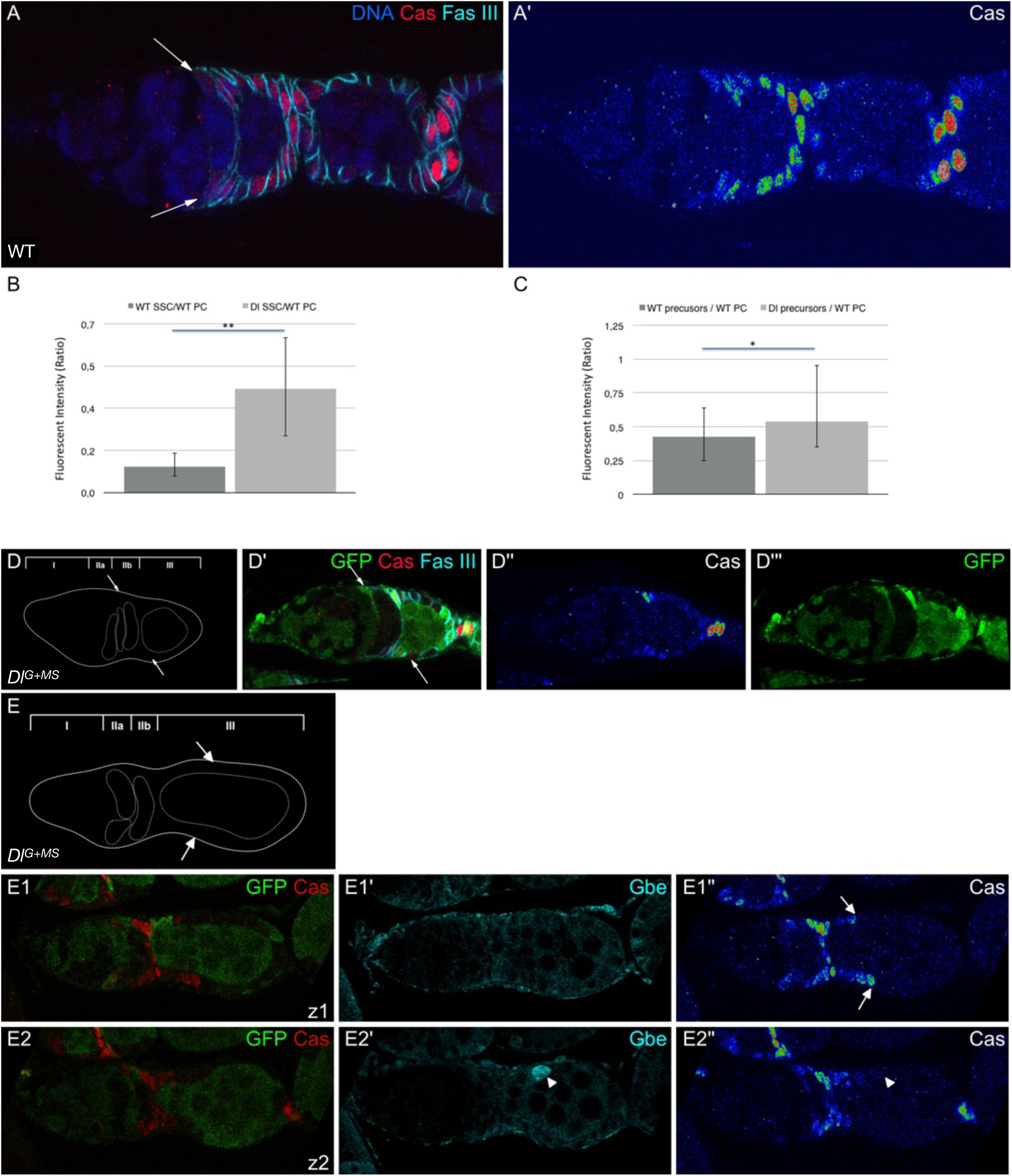
DlS controls Castor expression. The use of a color gradient allows the visualization of differences in intensities of accumulation from none (blue) to strongest (red) (A’, D’’, E1’’, E2’’). (A) WT germarium and stage 2 follicles showing the different levels of Cas expression in the ssc and pFC (weak level, arrows), the ppc-sc (intermediate level), the pc and the sc (high level). (B, C) Fluorescent intensity of Cas expression: ratio between the ssc and the pc (B) or between the precursors (mbfc or pc-sc) and the pc (C). Error bars indicate s.e.m. *, p<0.05; **, p<0.01 t-test versus control. (D) *Dl^G+MS^* germarium. A *Dl* cyst is present in region IIb. No Cas-positive cells are detected in regions IIa and IIb. (E) *Dl^G+MS^* germarium through 2 consecutive z-confocal sections (z1 to z2). A *Dl* cyst is present in region IIb. Cas-positive cells are detected in region III in *Dl* mbfc close to IIb (arrows). A *Dl* mbfc (visible in z2) in contact to WT mbfc, expresses high levels of *GbeSu(H)m8* (arrowhead), but does not express Cas.

**Figure S5.**
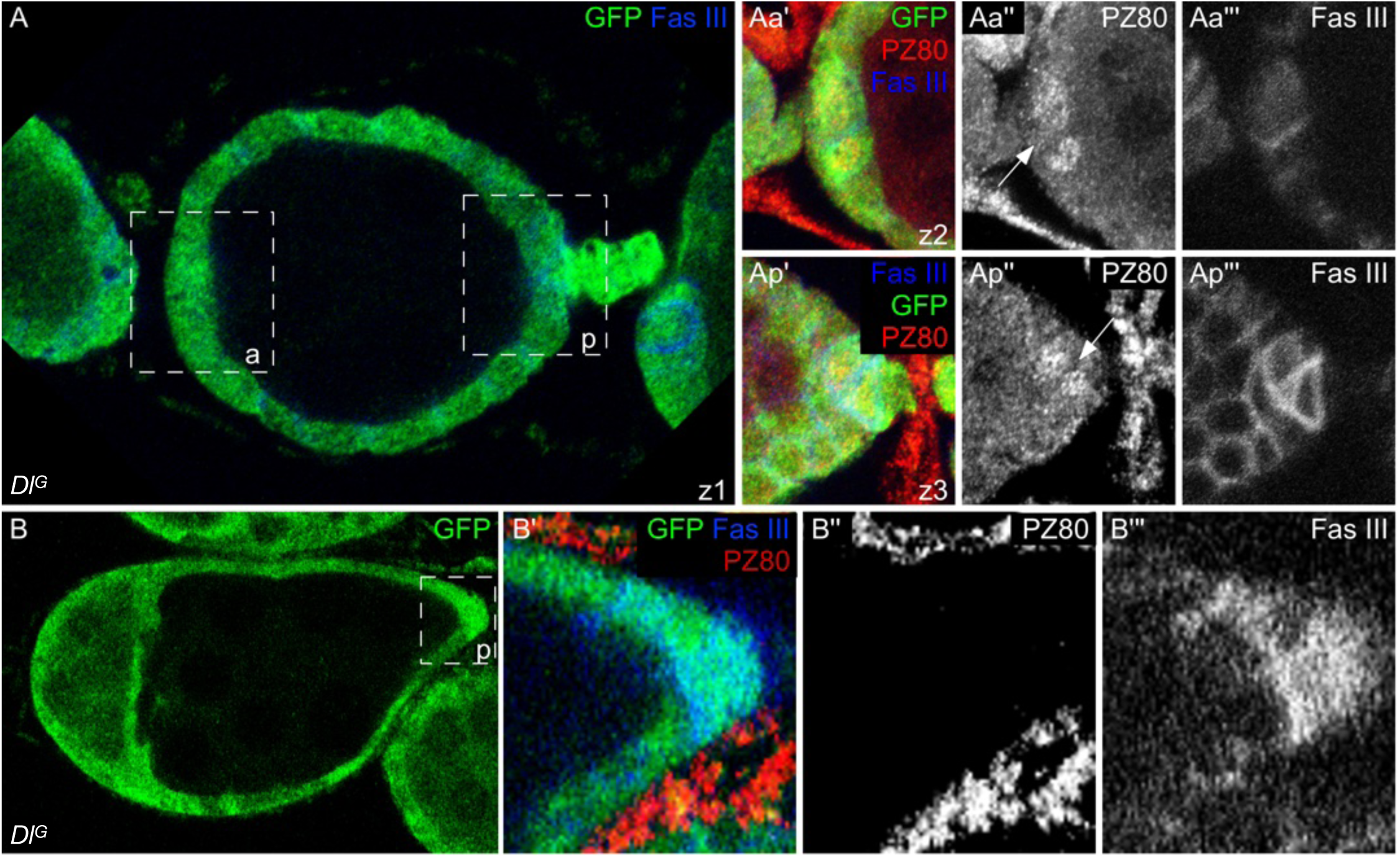
Robust polar cell differentiation requires DlG and DlS. (A-B) *Dl*^G^ follicles from females carrying the PZ80 enhancer trap. (A) Stage 3 follicle at 3 consecutive z-confocal sections (z1 to z3). Aa’ - Aa’’’ and Ap’ - Ap’’’ are magnified views of the anterior (a) and posterior (p) boxed areas in A. The arrows point to polar cells expressing PZ80. (B) Fused follicles between an anterior follicle with a WT cyst and a posterior follicle with a *Dl* cyst. B’-B’’’ are magnified views of the posterior (p) boxed area in B. No posterior cells express PZ80.

